# Landscape of IGH germline genes of Chiroptera and the pattern of *Rhinolophus affinis* bat IGH CDR3 repertoire

**DOI:** 10.1101/2023.01.20.524863

**Authors:** Long Ma, Longyu Liu, Jun Li, Hao Zhou, Jiaping Xiao, Qingqing Ma, Xinsheng Yao

## Abstract

The emergence and re-emergence of a number of viruses from bats that impact human and animal health has resulted in a resurgence of interest in bat immunology. Characterizing the immune receptor repertoire is critical to understanding how bats coexist with viruses in the absence of disease and developing new therapeutics to target viruses in humans and susceptible livestock. We annotated IGH germline genes of *Rhinolophus ferrumequinum* (RF), *Phyllostomus discolor* (RD) and *Pipistrellus pipistrellus* (PP), and investigated the evolutionary relationship between bat germline genes and that of human, mouse, cow, and dog. The IGH repertoire characteristics of *Rhinolophus affinis* bat (RA) were also analyzed. The V gene families of all three bat species can be classified into three Clan, although PD is special with the abnormal length of IGH locus and 22 reverse V genes. Moreover, the bats germline genes are quite differed from those of human, mouse, cow, and dog in evolution, but the three bat species have high homology. The CDR3 repertoire of RA are unique in many aspects including CDR3 subclass, V/J genes access and pairing, CDR3 clones and somatic high-frequency mutation compared with that of human and mouse, which may be the immunologic basis for the asymptomatic nature of viral infection in bats. This study provide immune genome information and extensive reference for the basic research of bat and virus infection mechanism.

## Introduction

Bats are the second largest mammal after rodents, make up more than 20% of extant mammals. Bats coexist with viruses in the absence of disease, which is divergence from primates and rodents. The discovery of bats carrying viruses can be traced back to the middle of the last century, such as Newcastle disease virus found in 1950^1^and Tacaribe virus found in 1963^2^. Now, More than 100 viruses have been detected or isolated from bats^3^, including many viruses that infect humans, such as hepaciviruses, pegiviruses^4^, influenza A virus^5^, hantavirus^6^, mumps and respiratory syncytial virus^7^, severe acute respiratory syndrome coronavirus-like virus^8,9^, MERS and severe acute respiratory syndrome coronavirus-2^10,11^. Studying the mechanisms of immune tolerance in bats could lead to new approaches to improving human health^12^.

Bats carry highly pathogenic viruses without symptoms, which should be attributed to their special innate and adaptive immune responses. The composition and function of Toll like receptor^13,14^, interferon^15^ and a variety of innate immune response genes have been preliminary elaborated in bats, and the mechanism of interferon in bats and in humans is different^16^, suggesting that bats have a stronger innate antiviral response and can control viral replication early^17,18^. However, it is not clear what role the adaptive immune response of bats plays in this process.

Revealing the mechanism of B cell response and antibody production in bats will help to clarify the mechanism of asymptomatic bats carrying viruses. In 1982, IgM, IgG and IgA were isolated from the serum of Artibeus lituratus and P.giganteus, and which were homologous with that of human immunoglobulin^19^. In 2010, the representative immunoglobulin heavy chain variable region (VH) genes of Pteropus Alecto and Pteropus vampyrus antibodies were found, involving all three VH families (I, II and III)^20^. In 2011, John et al. found the transcriptomic evidence of IgM, IgE, IgA and IgG subclasses in Chiroptera^21^, and bats showed high diversity of VH, DH and JH genes^22^. In 2021, Peter et al. annotated 66 IGHV genes, 8 IGHD and 9 IGHJ genes at the IGH locus of Egyptian rousette bats using bacterial artificial chromosome^23^. Although these previous studies provide a basis for understanding the humoral immune response of bats, to further explore bat B cell-mediated adaptive immune response depends on the annotation and application of bat IG germline genes at the chromosome level.

With the completion of genome sequencing and chromosome assembly in a few bats, Rhinolophus ferrumequinum (RF), Rousetus aegyptiacus, Phyllostomus discolor (PD), Myotis myotis, Pipistrellus pipistrellus (PP) and Molossus molossus^24^, we have finished annotation and preliminary application of the TR in RF^25^. Now, we unveiled a detailed map of Chiroptera IGH germline genes on chromosome level, and provided the first immune receptor repertoire of bat.

## Results

### The structure of bat IGH loci

The IGH loci of RF, PD and PP are located on chromosome 6 (CM014231.1), 15 (CM014268.2) and 20 (LR862376.1), with lengths of 350kb, 2530kb and 740kb, respectively. A total of 41 IGHV genes, four IGHD genes and six IGHJ genes were identified in RF (Fig.1A). A total of 81 IGHV genes (including 22 reverse IGHV genes) (Table S1), 16 IGHD genes and seven IGHJ genes were identified in PD (Fig.1B). A total of 57 IGHV genes, seven IGHD genes and six IGHJ genes were identified in PP (Fig.1C). Moreover, the IGHC genes of four immunoglobulins (IgM, IgG, IgE and IgA) were found in all three bat species.

**Fig.1.**
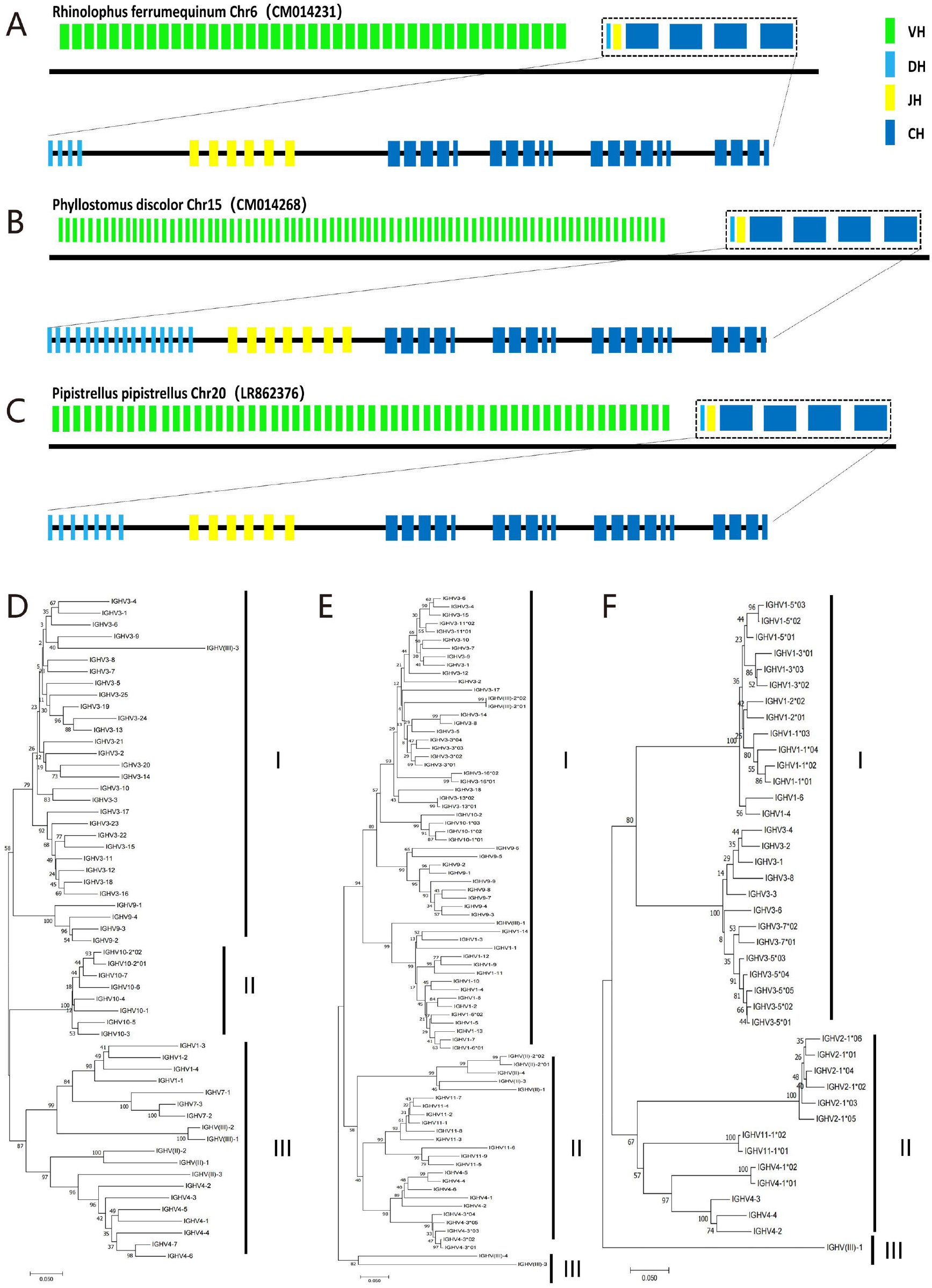
Structure of bat IGH loci and the three Clan of V genes. **A:** The IGH locus of Rhinolophus ferrumequinum. **B:** The IGH locus of Phyllostomus discolor. **C:** The IGH locus of Pipistrellus pipistrellus. **D:** The phylogenetic tree of Pipistrellus pipistrellus. **E:** The phylogenetic tree of Phyllostomus discolor. **F:** The phylogenetic tree of Rhinolophus ferrumequinum. Green segment is IGHV gene; light blue segment is IGHD gene; yellow segment is IGHJ gene; dark blue segment is IGHC gene.

### Nomenclature and amino acid composition of IGHV gene

The amino acid structures of all V genes of RF (Fig. S1A), PD (Fig. S1B) and PP (Fig. S1C) have classical conservative sites in the three skeleton regions, such as Cys23, Trp41 and Cys104, and the nucleotide sequence similarity is high, which are 39.7% - 99.3% (RF), 5.0% - 100.0% (PD) and 43.5% - 96.3%(PP), respectively.

IGHV genes of three bat species are classified and named (Table S2). The 41 IGHV genes of RF are divided into six gene families, of which only one pseudogene is a monogenic family, and the other five are polygenic families. The 81 IGHV genes of PD and the 57 IGHV genes of PP are divided into eight polygenic families respectively, and each contains two pseudogene families. The number of non functional V genes of the three bat species are six (RF), 33 (PD) and 23 (PP), respectively (Table S3).

Comparing the bat IGHV genes with more than 20 species in the IMGT database, bats have great genetic differences with other species in the number of IGHV families, functional genes, pseudogenes, ORFs, and the composition of polygenic families. However, as shown in the phylogenetic tree (Fig.1D-F), the IGHV genes of the three bats can be divided into three Clans: ClanI, ClanII and ClanIII, which are similar to those of mammals such as human and mouse. We further analyzed the evolutionary relationship of V genes between bat and human, pike, cow, and found no species with significant convergence in bats. Compared with carnivores, primates and artiodactyls, bats show uniqueness in the number of IGHV genes (Fig. S2).

### Nomenclature and amino acid composition of IGHJ gene

Six, seven, and six IGHJ genes were identified in IGH loci of RF, PD and PP, respectively, and all of them are functional genes. The amino acid sequence alignment showed that the number and characteristics of the IGHJ genes of the three bat species are consistent with those of human beings. The J genes of the three bats have conserved WGQG and VTVS structures except for one amino acid changed in IGHJ2 and IGHJ4 of the PP (Fig.2A).

**Fig.2.**
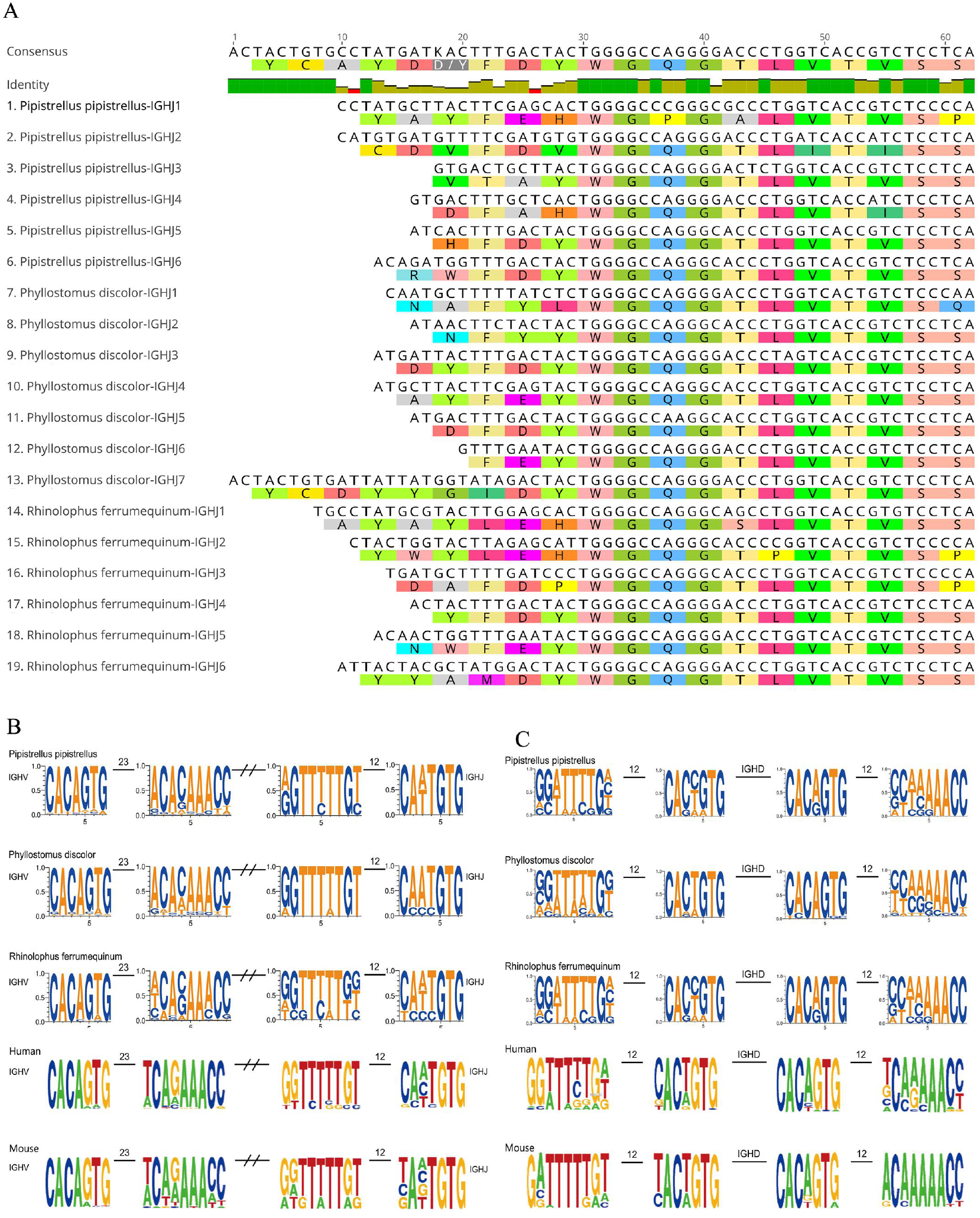
The structure of IGHJ genes and RSS sequence. **A:** Sequence comparison of all IGHJ genes in three bat species. **B:** RSS characteristics of V and J genes in bat, human and mouse. **C:** RSS characteristics of D genes in bat, human and mouse.

### The characteristics of bat 12/23 RSS

23 RSS are located behind the IGHV genes and each RSS contains a heptamer and a ninomer. All the heptamer sequence of three bat species are relatively conservative (Fig.2B), while the nucleotides at the fourth position of the ninomer are diverse. 12 RSS were located on front of IGHJ genes and before and after IGHD genes, and each RSS contains a heptamer and a ninomer. The GTG nucleotides at the last three positions of heptamer and the TTTTT nucleotides at positions 3-7 of ninomer in the IGHJ pre-12 RSS sequence of the three bats are relatively conservative. For the pre-12 RSS of IGHD genes, the relatively conservative nucleotide sites are G at position 8 in the ninomer, CA at positions 1 and 2 and TG at positions 6 and 7 in the heptamer in the three bat species (Fig.2C). These conservative sites have not undergone any mutations in the three bats. The conserved nucleotide of the post-12 RSS of IGHD genes in the three bat species are the third C in heptamer and the last four AACC in the ninomer.

### Construction of RA IGH CDR3 repertoire

As expected, the construction of IGHM, IGHG, IGHA and IGHE showed obvious peaks at the 1500bp, suggesting that the constructions were successful. Although the number of total unique IGH CDR3 sequence and each subclass CDR3 sequence are divergence among three RA (Table 1), they all meet the CDR3 sequence analysis requirements (the unique clone sequence / total functional sequence < 10%). Moreover, quality control was performed on the IgG of each bat by designing two sets of primers. The number of sequences obtained was identical, and the common high-frequency sequences proportion in each sample reached more than 52% (Fig. S3). Interestingly, the sequence composition of RA IGH are high homology with that of the annotated RF, and only a few V and J gene families are homologous with that of the annotated PD and PP (Fig.3).

**Table 1.** The sequencing statistics of Rhinolophus affinis IGH CDR3.

| Sample | IGH |  | Subclasses |  |  |  |
| --- | --- | --- | --- | --- | --- | --- |
|  | Productive | Clonotype | Name | Productive | Clonotype | Clonotype/Productive |
| B1 | 1,611,347 | 14,694 | IGG | 1,600,169 | 13,865 | 0.87% |
|  |  |  | IGA | 7,742 | 420 | 5.42% |
|  |  |  | IGM | 3,173 | 372 | 11.72% |
|  |  |  | IGE | 263 | 37 | 14.07% |
| B2 | 4,240,438 | 32,130 | IGG | 2,159,687 | 21,556 | 1.00% |
|  |  |  | IGA | 302,272 | 503 | 0.17% |
|  |  |  | IGM | 1,747,415 | 9,797 | 0.56% |
|  |  |  | IGE | 30,974 | 275 | 0.89% |
| B3 | 11,270,098 | 59,574 | IGG | 10,517,634 | 49,456 | 0.47% |
|  |  |  | IGA | 238,894 | 668 | 0.28% |
|  |  |  | IGM | 506,333 | 9,307 | 1.84% |
|  |  |  | IGE | 7,237 | 143 | 1.98% |

B1: bat 1; B2: bat2; B3: bat3.

**Fig.3.**
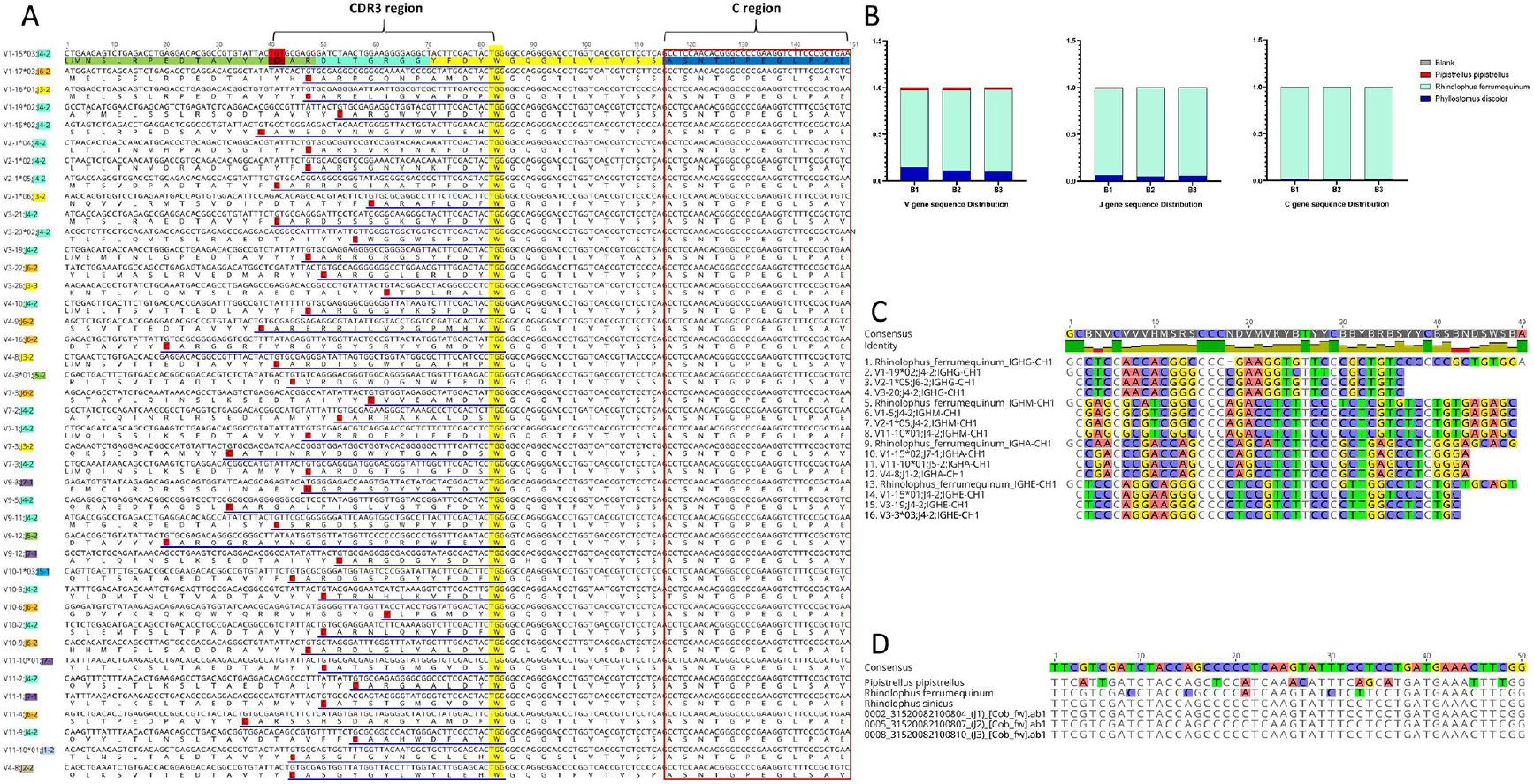
Alignment of Rhinolophus affinis sequence with that of annotated three bat species germline genes. **A:** Partial sequences of Rhinolophus affinis. **B:** The proportion of V, J and C genes of three annotated bats in the Rhinolophus affinis. **C:** The comparison of C region between Rhinolophus ferrumequinum and Rhinolophus affinis. **D:** Sequence comparison of Cytb gene in four bats. All the sequences of Rhinolophus affinis were obtained by sequencing.

### The comparison of RA IGH CDR3 subclass

The IGH CDR3 sequences of human and mouse were also included in this study to further contrastive analysis of the characteristics of IGH CDR3 between bats and other species (Table S4). The IGH CDR3 subclass with the largest number of sequences of RA is IgG, followed by IgM, IgA and IgE, while IgM is the most at the transcriptome level in human and mice, followed by IgA and IgG, which is consistent with published articles and shared database obtained from healthy people and mice.

### V/J access and pairing of IGH CDR3

The V and J access of IGH CDR3 in RA, humans and mice are shown in Fig.4A. The three species are biased towards IGHV1. Bats and people also favor IGHV4, and IGHV3 is only available in bats at high frequencies. The frequency of other IGHV gene families in the three species is very low. IGHJ4 appeared frequently in all three species, and the frequency of IGHJ family access in bats is almost the same as that in humans, in descending order of access frequency, are: IGHJ4, IGHJ6, IGHJ5, IGHJ3, IGHJ1 and IGHJ2. Mice have no preference for the IGHJ families. Moreover, the access trend of V and J is consistent with shared data (Fig. S4A), and the V and J utilization of IGH subclass (IGM and IGE) are also similar to the overall (Fig. S5A-B). V/J pairing of RA is identical to that of human and mouse (Fig.4B-D; Fig. S4B, Fig. S6), which IGHV1-IGHJ4 pairing with high frequency was detected.

**Fig.4.**
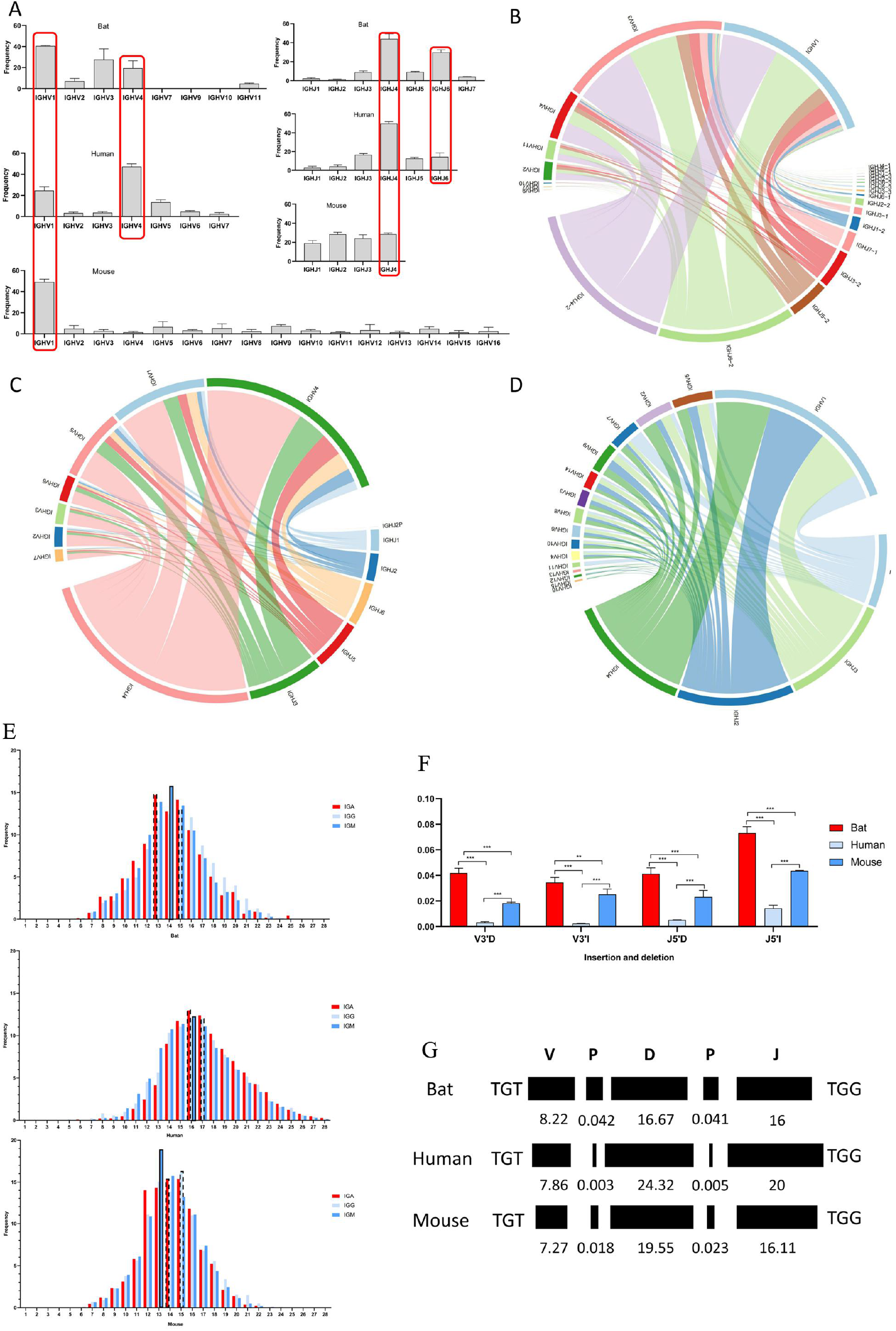
Analysis of Rhinolophus affinis IGH CDR3 repertoire. **A:** V/J gene access of Rhinolophus affinis, human and mouse. **B-D:** V-J pairing in Rhinolophus affinis, humans and mouse, respectively (only one sample of each species randomly selected for display, and the rest is shown in Supplement figure). **E:** CDR3 length distribution of IGH subclasses in Rhinolophus affinis, humans and mouse, respectively. **F:** The insertion and deletion of CDR3 region of Rhinolophus affinis, humans and mouse, respectively. **G:** The CDR3 length composition of Rhinolophus affinis, humans and mouse, respectively. ns: P>0.05; *:P<0.05; **: P<0.01; ***: P<0.001.

### Length distribution of IGH CDR3

The CDR3 length of IgA, IgG and IgM are bell shaped (Fig.4E). Bat is centered on 13,15,14 AA, mouse is centered on 15,15,13 AA, and human is centered on 16,17, 16 AA. The difference of the composition of V, D, and J (Fig.4F) and deletion /insertion (Fig.4G) of CDR3 regions in bats, humans, and mice were also carried out. The deletion and insertion of bats at V3’ and J5’ ends are higher than those of mice and humans. The length of the terminal of the V gene in the CDR3 region of bats is the longest, while the length of the D gene and the J gene are the shortest among the three species. In addition, the IgG showed a longer AA distribution than IgA and IgM in all three species. The length distribution and AA access of shared IGH CDR3 data also match the above results (Fig. S4C).

### The clones of IGH CDR3 repertoire

Fewer than 100 CDR3 clones were defined as rare clones, and the proportion of rare clones in bats is significantly lower than that in humans and mice (Fig.5A-B). The cloning frequencies of human and mouse are similar on the whole, while bats show high individual differences (Fig. S7A-B). The Shannon index indicate that the IGH CDR3 repertoire diversity of bats is lower than that of humans and mice, but there is no significant difference (Fig.5C).

**Fig.5.**
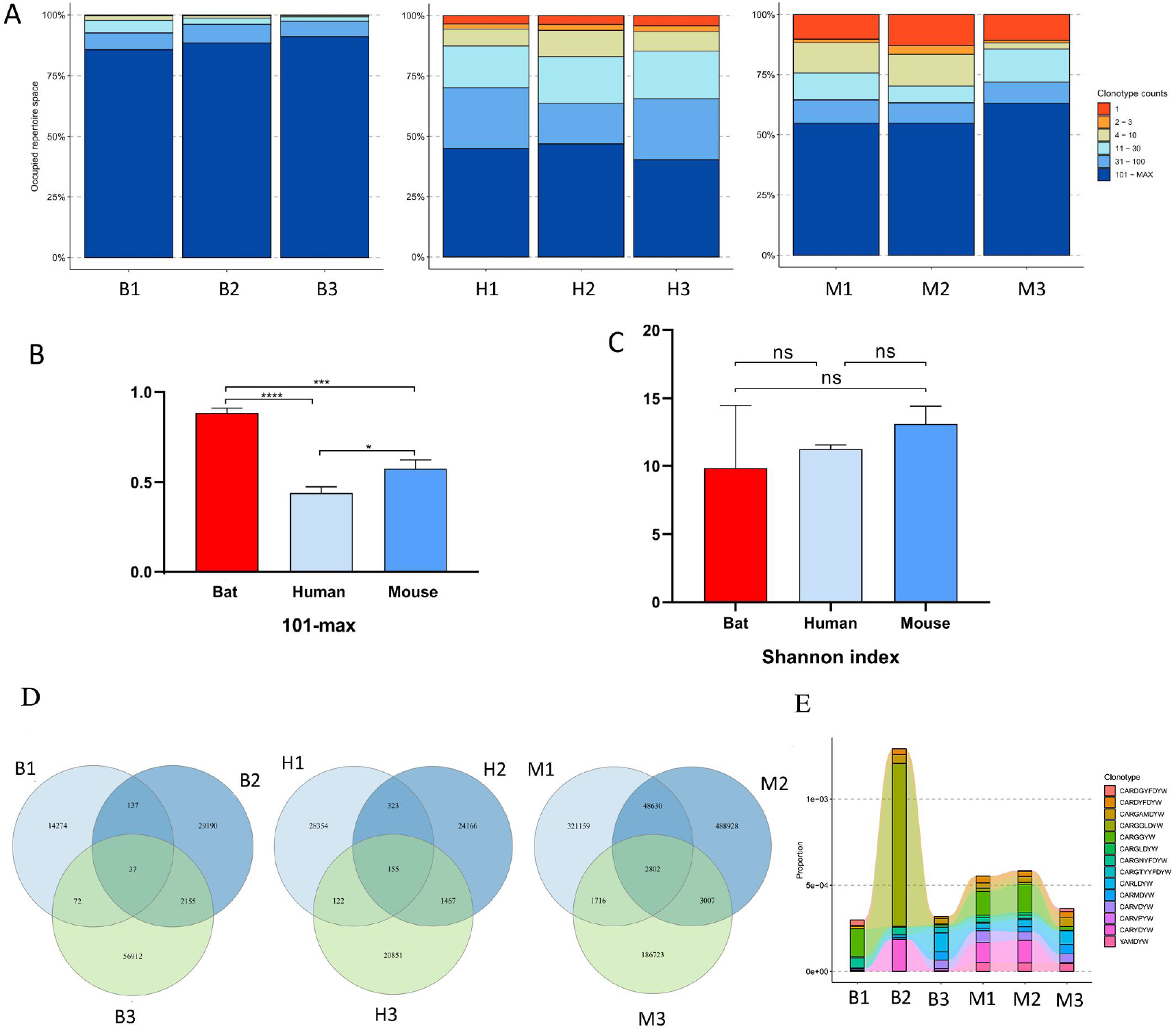
The clones analysis. **A:** Distribution of rare clones in Rhinolophus affinis, humans, and mouse, respectively. **B:** Statistics of clones above 100 in Rhinolophus affinis, humans and mouse. **C:** The Shannon index of Rhinolophus affinis, humans and mouse. **D:** The public clones of Rhinolophus affinis, humans and mouse, separately. **E:** The clonotype tracking between Rhinolophus affinis and mouse. B: bat; H: human; M: mouse. ns: P>0.05; *: P<0.05; **: P<0.01; ***: P<0.001.

The IGH CDR3 unique clones overlapped among bats is low, and the highest is that of mice (Fig.5D). For IG subclass, the IGG unique clones overlapped among humans is most, and it is inconsistent with the total IGH. The shared unique clones of IGA and IGM are similar with the total IGH in the three species (Fig. S7C-E). Notably, clonotype tracking showed 14 public clones between bats and mice (Fig.5E).

The AA intake of IG subclasses (IgG, IgA, IgM) in bats is highly consistent with that of humans and mice (Fig.6A), with high-frequency intake of Y, G, A, R, W, D. The IGH CDR3 motif showed specificity in the three species (Fig. S8), and we counted the top 10 motifs of each species and found that bats and mice have multiple same motifs with the high frequency (Fig.6B), such as YFDYW, AMDYW, YAMDY, YYFDY and YYAMD. There is only one intersection in the top 10 motifs of bats and humans, YFDYW. Moreover, the top 10 motifs of IG subclasses (IgG, IgA, IgM and IGE) were also analyzed. IgA, IgG and IgM have no significant special high-frequency motifs in the three species except that the order of motif frequency is slightly different (Fig. S9), while the IgE in bats and mice shows species differences (Fig. S10). We further analyzed the four subclasses of bats and found that IgA and IgE have multiple unique high-frequency moitf, while IgM and IgG do display almost the same high-frequency moitfs. Moreover, each analyzable sequence contains a complete J region. The SHM of bat J region is significantly higher than that of human and mouse (Fig.6C).

**Fig.6.**
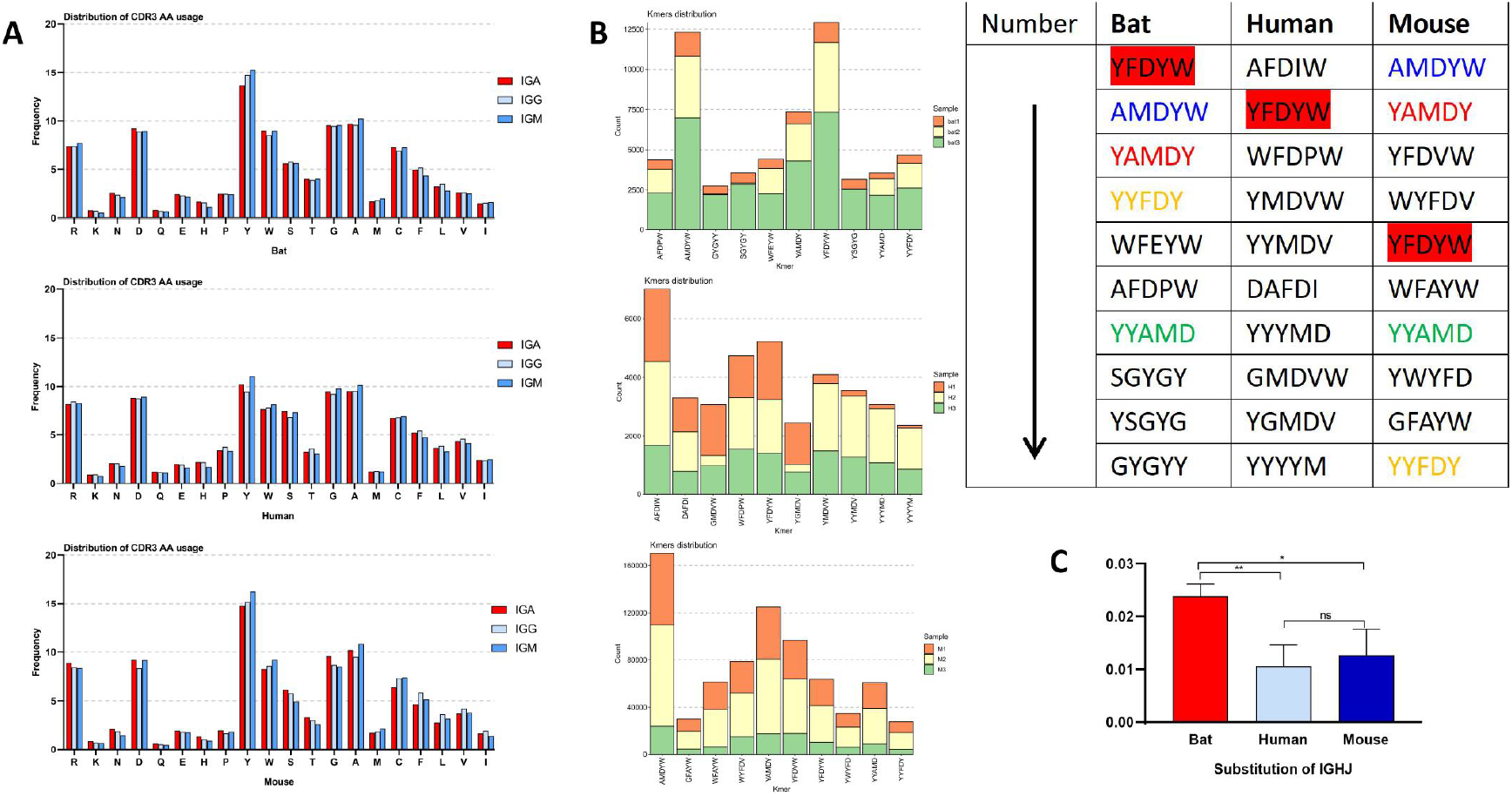
The AA usage and motif composition of CDR3 region and mutations in IGHJ region. **A:** Statistics of AA usage of IGH subclasses in Rhinolophus affinis, humans and mouse, respectively. **B:** The top 10 motif of IGH in Rhinolophus affinis, humans and mouse, respectively. **C:** Statistics of mutations in IGHJ region of Rhinolophus affinis, humans and mouse. B: bat; H: human; M: mouse. ns: P>0.05; *: P<0.05; **: P<0.01; ***: P<0.001.

## Discussion

Bats are carriers of highly pathogenic viruses, which have caused massive damage to human health and pose a huge risk to the spread of viral diseases in the future^26^. It is urgent to clarify the relationship between bat immune system and virus. Rabies virus^27^ and Australian bat lyssavirus^28^ can cause clinical symptoms in bats, suggesting that the immune system of bats is also struggling with the virus. We have finished the TR germline genes annotation of RF^25^. In this study, we annotated the IGH germline genes of three bat species completely and explored the characteristics of bat’s IGH CDR3 repertoire. Compared with bat T cell response, studying the mechanism of bat B cell response and antibody production will better elaborate the reason why bat coexist with virus. Early studies found that the intensity and duration of neutralizing antibody reaction of Eptesicus fuscus remained lower than that of cavia porcellus and rabbits^29^. The Artibeus jamaicensis experimentally infected with Venezuelan encephalitis virus produced strong neutralizing antibody, but the detectable antibody response of P. discolor was slower and of lower magnitude and shorter duration than that of Artibeus^30^. Neutralizing antibodies against Ebola virus^31^, Hendra virus and SARS-like coronavirus^8^ were also detected in wild bats. These studies suggest that the annotation of bat Ig and its application to the study of the mechanism of bat B cell response will play a vital role in clarifying bat specific immune response. The characteristics of BCR repertoire of human, mice and other mammals are available and have applied in basic research and clinical diagnosis, while the composition and diversity of BCR libraries of bats are almost unknown.

This study is the first time to completely annotated the IGH germline genes of RF, PD and PP. The length of IGH heavy chain in most mammals recorded by IMGT is similar, but there are obvious differences among the three bats (430kb, 2500kb and 350kb), which may be related to the long-distance distribution of IGHV genes in PD. The IGH germline genes of the three bats are highly homologous, and also have high homology with the Egyptian rousette bats recently annotated using bacterial artificial chromosome^23^. Comparing the germline genes of bats with those of human, mouse, cow and dog, bats did not show significant homology with one of the species, but the V, D, J genes on the IGH chain of the three bats showed a clustered arrangement of similar genes. Interestingly, there are 22 reverse IGHV in front of the D/J genes in the PD, which rarely occurs in the annotated IG and even TR genes. This abnormality of the PD and whether there is a similar arrangement in other bat species will be of great concern.

M. lucifugus, E. fuscus, C. perspicillata and C. Sphinx have 73, 20, 16 and 15 IGHV genes, respectively^21^. In this study, 57, 41, and 81 IGHV genes were identified in PP, RF and PD, respectively, and were different from Egyptian rousette bats^23^. The difference in the number of V genes in different bat species may suggest that bats with many species are different from other mammals, and may have the significant diffusion of IGHV gene. We mapped the evolutionary tree of IGHV genes of three bats, which is consistent with the IGHV family of mammals such as human and mouse, and can be classified into Clan I, II and III, suggesting that the evolution of bats IGHV genes is consistent with that of human and mouse.

Among the various species shared by IMGT, the proportion of IGHV pseudogenes are high, 63.3%, 35% and 41.6% in human, pig and mouse, respectively. In this study, 31.6%, 39.5% and 12.2% pseudogenes were found in PP, PD and RF, respectively. Pseudogenes are slightly lower in bats but do not show significantly difference compared to other species, suggesting that there is no significant difference in the process of IGHV gene becoming pseudogene by random and extensive mutation in different species.

The length of IGHJ gens are generally 37-63bp among the species recorded in the IMGT database. In this study, The length of IGHJ gene of the three bat species are 45-65bp, suggesting that bats are similar to other species in length. We analyzed the IGHV and IGHJ sequences in three bat species and found that Cys23, Trp41 and Cys104 were highly conserved in the IGHV sequences, and IGHJ gene has highly identical amino acid conservative sequences. In a total of 19 IGHJ genes, except for two genes which have one mutation, the other IGHJ sequences have WGQG and VTVS structures, which are basically consistent with that of other mammals recorded in the IMGT database, suggesting that the IGH genes of bats conforms to the standard pattern of real mammals, but is different from birds or protomammals.

RSS is one of the most critical components of adaptive immune evolution. RSS before and after V and J genes found in this study are classic RSS, and the conservative sites of RSS sequences of the three bats are similar, whether it is 7-mer or 9-mer. Compared to human and mouse, the conserved positions of nucleotides are basically the same, suggesting that the RSS sequences of bat annotated in this study are consistent with the classic RSS sequences of mammals, and have not changed greatly in the process of species evolution. However, whether bats have nonclassical RSS, such as spacer 12 ± 1 bp or 23 ± 1 bp, remains to be further explored.

Some bat species may lose IgD isotype in evolution. The transcribe genes encoding IgA, IgG, IgM and IgE subclasses were found in Cynopterus sphinx, Carollia perpicillata, M.lucifugus, E.fuscus and two short-nosed fruit bat, but the IgD transcripts were only recovered from insectivorous bats and were comprised of CH1, CH3 and two hinge exons^21^. Moreover, no transcripts of IgD were detected in P.alecto^17^. In this study, We analyzed the characteristics of the C-region of the IGH chain in RF, PD and PP, similarly, there are four C-regions: IGHM, IGHG, IGHE and IGHA, but no IGHD.

Many evidences show that different bat species have common ancestors in the evolution(*18, 26, 32*). According to the high homology of the annotated IGH in RF, PD and PP, we have established a method to analysis RA IGH CDR3 repertoire, and made a preliminary comparison with that of human and mouse.

The sequencing for bat IGH CDR3 repertoire was successful. According to the IGH CDR3 sequence and isotypes composition of RA, we found a high homologous between RA and RF, suggesting that the adaptive immune response of bats can be studied at the level of ‘family’. There are 18 families in bats, and the evolution of B cell IGH loci are quite different. The study of adaptive immune responses in bats may be more complex than other mammals. The IGH CDR3 isotype of bats differ significantly at the trahscriptome level from human and mouse, such as the extremely low proportion of IgA. Moreover, the serum IgA of healthy bats was significantly lower than expected, suggesting that higher quantities of IgG in mucosal secretions may be compensation for this low abundance or lack of IgA^33^. The characteristics of bats that differ from humans and mice in isotypes may be the reason for the bat special immune response.

The length of IGH CDR3 in each species is mainly caused by the differences of IGHV terminal, IGHD, IGHJ front-end, and insertions and deletions in the rearrangement. In general, the length of IGH CDR3 is positively correlated with body size, and B cells may also change CDR3 length during self tolerance selection. In our previous study, we found that the shear of V3’ and J5’ ends of human in IGH CDR3 is higher than that of mouse^34^. However, the results of this study are opposite, and the consistency of bat and mouse is higher than that of human, suggested that the insertion and deletion of IGH may be more frequent in the development and tolerance of B cells in bat and mouse species than in human beings.

The clonality and diversity of IGH CDR3 repertoire are related to the intensity and breadth of adaptive immune response. The abundance of rare clones in RA is low, and the ultra-high clones is high, which showed obvious differences with human and mouse. The possible reason is that bats show a strong B-cell response to a small number of antigens, but the breadth of response to antigens is lower than that of humans and mice. Certainly, small samples size is a defect of this study, and a consistent sequencing depth is also needed to further explore the diversity of IGH CDR3 repertoire in bats.

In general, there are abundant common T cell clones and B cell clones between different individuals of human or mouse^35^. However, the source and effect of these public clones are not clear at present. Among the three species in this study, mice have the highest rate of public clones, which may be consistent with the genetic background of Balb/c mice. Notably, bats and mice share some clones, suggesting that the characteristics of overlapping clones can be used to explore the response differences between bats and other species.

The AA uptake in IGH CDR3 region was highly consistent among bat, human and mouse in this study, suggesting that the IGH CDR3 region in mammals has great commonality in composition, structure and antigenic determinant combination. In P. lecto, the IGHV region was rich in Arg and Ala, and the amount of Tyr is small, which may lead to the evolution of antibodies in bats, whose polymerizing reactivity is low and only weakly associated with antigens^20^. However, the Tyr content of the three RA in this study is almost the same as that of human and mouse.

The composition and conformation of motif in CDR3 region are important for B/T cells binding to the corresponding antigen. Among the top 10 high frequency motifs in IGH CDR3, bats and mice showed higher similarity, with 5 same motifs. The high frequency motifs of each subclass are basically consistent with the total IG, but bat IGE and mouse IGE showed less common high frequency motifs than other subclasses. Bat IGA and IGE showed multiple unique high frequency moitfs, while IGM and IGG show almost the same high frequency moitfs, revealed that the classification conversion mechanism of IgE and IgA in bats is inconsistent.

Different bat species may have different SHM. In black flying fox^20^ and fruit-eating bat^36^, only few SHM were found, while significantly more mutations were detected in RA IGH J sequences compared to human and mouse in this study. Bat’s higher SHM may be closely related to its tolerance or response to the virus, which is a breakthrough point to further explore the mechanism of bat B cell response and whether it produces high affinity antibodies to respond to the virus through SHM.

Many studies on bats’ high heart rate and metabolism, long life span, low tumor incidence, asymptomatic ability to carry and transmit highly pathogenic viruses have been carried out, including the cell lines establishment of pteropid bat^37^, the preparation of polyclonal antibodies of bat IgG, IgM and IgA^33^, the sequencing and assembly of bat genome^24^, and the establishment of the Bat1K genome consortium unites^38^, etc. These efforts will provide the basis and technology for elucidating the innate and adaptive immune responses of bats. This study displayed the IGH germline genes of three bat species at the chromosome level and analyzed the characteristics of bat IGH CDR3 repertoire, which provided a new technology and basic data for studying the IGH characteristics and the mechanism of antiviral immune response in bat.

## Materials And Methods

### Location of V, D and J genes of IGH locus

The whole genome sequence information of RF (GCA_00415265.2), PD (GCA_004126475.3) and PP (GCA_903992545.1) were obtained from NCBI website (https://www.ncbi.nlm.nih.gov/). The classic IMGT_ LIGMotif ^39^ and 12/23RSS^40^ approach were adopted to identify bat’s IGH germline genes. The chromosomal location of IGH loci were determined by comparing mammals IGHC genes that are available on IMGT website (https://www.imgt.org/genedb/) with the whole genome sequence of three bats species. Similarly, mammalian IGHV, IGHD, and IGHJ sequences download from IMGT website were mapped with the chromosomes determined by bat’s IGHC gene to obtain possible germline genes, and these genes were labeled with Geneious Prime software. Next, the sequences from IGHV to IGHC were selected with 10KB as a group, and were dropped into Ig BLAST website (http://www.ncbI.nlm.nih.gov/igblast/) to screen possible germline genes. Moreover, Meme website (http://meme-suite.org/) was applied to screen the possible RSS sequences for finding unlabeled IGHV, IGHD and IGHJ genes, and verifying all germline genes. Finally, the genes with complete initiator codon, splice site, sequence length greater than 271bp and proper RSS at the end of the sequence were identified as IGHV genes.

### Characteristic analysis and nomenclature of bat germline genes

The characteristics of each germline gene of the three bat species were analyzed according to IMGT guidelines^41^. The labeled V, D, and J gene sequences were uploaded to IMGT/V-Quest (http://www.imgt.org/IMGT_vquest/input), and were defined as functional genes (F), open reading frame (ORF) and Pseudogene (P) according to IMGT functional classification principles. The similarity of amino acids and nucleotides of three bat’s germline genes were performed in Geneious Prime software, and the conserved amino acid, CDR and FR regions of the V and J genes in three bat species were also labeled using Geneious Prime software and IMGT/V-Quest.

According to the naming rules of human IGH in IMGT, those with nucleotide similarity ≥75% were classified into the same family, the IGHV genes of three bats were clustered with human IGHV gene families, and those with nucleotide similarity ≥75% were classified into the same family, which was named uniformly with that of human. Phylogenetic trees of IGHV/J genes of three bats species were also constructed respectively in MEGA version 7 by the neighbor-joining method. The Logo graph was drawn to analyze the composition characteristics and conservative of RSS using Weblogo website (https://weblogo.threeplusone.com/create.cgi).

### Construction of bat IGH reference dataset

Due to the high homology of IGH V, D, J and C sequences of the annotated three bat species, the bat IGH reference gene bank was constructed for the first time by using the annotated IGH germline genes of the three bat species, with a total of 179 IGHV genes (including 57 pseudogenes), 27 IGHD genes, 19 IGHJ genes and 12 IGHC genes. These V and J genes that have been included in the reference dataset were divided into four skeleton regions (FR1, FR2, FR3, FR4) and three variable regions (CDR1, CDR2, CDR3) according to the amino acid conservative sites which been recognized by MiXCR^42^. Finally, bat IGH reference dataset was constructed and used for bat IGH repertoire analysis.

### Sample preparation and IGH CDR3 repertoire sequencing

The muscle and spleen of bats were collected in Zunyi, Guizhou Province, China. The muscle tissues were used to extract genomic DNA, and the Cytb gene was amplified to determine the genotype of bats (Table S5). In this experiment, three Rhinolophus affinis were selected for the IGH CDR3 repertoire construction and characteristic analysis. The spleen tissues were used to extract total RNA. The construction and sequencing of the library were conducted by Hangzhou ImmuQuad Biotechnologies Ltd using the 5 ‘RACE method. The primers were designed in the conservative region, which was obtained by comparing all available bat’s IGHC genes (Figure S11). In order to control the quality of library construction, two groups of primers with different specificity were designed for IGG, and the efficiency of high-frequency sequence amplification was analyzed between the two groups. After quality control of sequencing raw data, MiXCR software was applied for subsequent analysis using the bat IGH reference dataset we created.

We also compared the characteristics of bat IGH CDR3 repertoire with those of human and mouse. Peripheral blood of three healthy volunteers (male, 19-25 years old) and bone marrow of three mice (two months old) were collected for construction and sequencing of IGH CDR3 repertoire. Moreover, We also downloaded the public IGH CDR3 repertoire data for further comparative analysis (including human and mouse), both of which were obtained by 5 ‘RACE method. This study was approved by the animal protection and ethics committee of Zunyi Medical University.

### Analysis of the IGH CDR3 repertoire

The MiXCR, VDJtools and immunarch software were used to analyze the composition of each BCR CDR3 sequence, including: nucleotide, amino acid (AA), count (reads), frequencyCount (%), CDR3 Length, the V-J rearrangement of the CDR3 repertoire, the proportion and frequency of unique CDR3 sequences, CDR3 repertoire clonality, CDR3 amino acid length, CDR3 animo acid usage, V deletion and J deletion, and dominant V-J combination gene segments were also calculated in different samples. To assess the clone frequency of CDR3 region, the inverse Shannon index was performed.

### Statistical analysis and graphing

R package “ggplot2”, “Venn Diagram”, and GraphPad Prism (version 5) were used to plot the figures. Data analysis was performed by R studio (v3.3.3) and GraphPad Prism (version 5) software. P-values were calculated with the aid of the t test. P<0.05 was considered statistically significant.

## Supporting information

BioRXIV

## Acknowledgments

We would like to express our gratitude to Jiang Zhou and Xingliang Wang of Guizhou Normal University for their help in collecting wild bats, and Hangzhou ImmuQuad Biotechnologies Ltd for bat IGH CDR3 library building and high-throughput sequencing.

## Author contributions

XY and LM designed the research, LL and JL did the experiment and wrote the paper. HZ, JX and QM analyzed parts of the data. All authors contributed to the article and approved the submitted version.

## Data and materials availability

The original data sets generated and analyzed during this study are made available by the corresponding author upon reasonable request.

