## Supplementary material for "Landscape of IGH germline genes of Chiroptera and the pattern of *Rhinolophus affinis* bat IGH CDR3 repertoire": BioRXIV

Table S1 22 reverse IGHV genes in Phyllostomus discolor.

| Name | Minimum | Maximum | Length | Direction |
|---|---|---|---|---|
| IGHV(II)-2*02 | 2057623 | 2057921 | 299 | Reverse |
| IGHV(III)-1 | 2661200 | 2661495 | 296 | Reverse |
| IGHV(III)-2*01 | 2627687 | 2627979 | 293 | Reverse |
| IGHV1-1 | 2671585 | 2671876 | 292 | Reverse |
| IGHV1-2 | 2618909 | 2619204 | 296 | Reverse |
| IGHV1-7 | 979077 | 979372 | 296 | Reverse |
| IGHV3-1 | 2706384 | 2706682 | 299 | Reverse |
| IGHV3-2 | 2674087 | 2674379 | 293 | Reverse |
| IGHV3-3*01 | 2638006 | 2638301 | 296 | Reverse |
| IGHV3-4 | 2513676 | 2513968 | 293 | Reverse |
| IGHV3-5 | 2489863 | 2490155 | 293 | Reverse |
| IGHV3-11*02 | 2025107 | 2025402 | 296 | Reverse |
| IGHV3-13*01 | 2038667 | 2038964 | 298 | Reverse |
| IGHV3-14 | 1961559 | 1961851 | 293 | Reverse |
| IGHV4-1 | 2540331 | 2540629 | 299 | Reverse |
| IGHV4-2 | 2533135 | 2533430 | 296 | Reverse |
| IGHV4-3*04 | 2049536 | 2049831 | 296 | Reverse |
| IGHV9-1 | 2579768 | 2580069 | 302 | Reverse |
| IGHV9-3 | 965030 | 965332 | 303 | Reverse |
| IGHV10-1*01 | 2600109 | 2600407 | 299 | Reverse |
| IGHV11-1 | 2500176 | 2500468 | 293 | Reverse |
| IGHV11-3 | 1348145 | 1348435 | 291 | Reverse |

Table S2 Statistics on the number, direction and functionality of genes contained in the IGHV gene family of three species of bats.

| IGHV subgroup | Rhinolophus ferrumequinum | | Phyllostomus discolor | | Pipistrellus pipistrellus | |
|---|---|---|---|---|---|---|
| | direction | total | direction | total | direction | total |
| IGHV1 | + | 14 | +/- | 15 | + | 4 |
| IGHV2 | + | 6 | + | 0 | + | 0 |
| IGHV3 | + | 13 | +/- | 24 | + | 25 |
| IGHV4 | + | 5 | +/- | 10 | + | 7 |
| IGHV5 | + | 0 | + | 0 | + | 0 |
| IGHV6 | + | 0 | + | 0 | + | 0 |
| IGHV7 | + | 0 | + | 0 | + | 3 |
| IGHV8 | + | 0 | + | 0 | + | 0 |
| IGHV9 | + | 0 | +/- | 9 | + | 4 |
| IGHV10 | + | 0 | +/- | 4 | + | 8 |
| IGHV11 | + | 2 | +/- | 9 | + | 0 |
| IGHV(I) | + | 0 | + | 0 | + | 0 |
| IGHV(II) | + | 0 | +/- | 5 | + | 3 |
| IGHV(III) | + | 1 | +/- | 5 | + | 3 |
| Total | + | 41 | +/- | 81 | + | 57 |
| Functional | + | 35 | +/- | 48 | + | 34 |
| ORF | + | 1 | + | 1 | + | 3 |
| Pseudogene (P) | + | 5 | +/- | 32 | + | 20 |
| P/Proportion | + | 12% | +/- | 40% | + | 35% |

+ is forward;- is reverse; + /- exists in both forward and reverse directions.

Table S3 Functional determination of IGHV/IGHJ genes in three species of bats.

| IGHV/ IGHJ | Pipistrellus pipistrellus | | | IGHV/IGHJ | Rhinolophus ferrumequinum | | | IGHV/IGHJ | Phyllostomus discolor | | |
|---|---|---|---|---|---|---|---|---|---|---|---|
| | Functionality | Stop Codon | Defective RSS | | Functionality | Stop Codon | Defective RSS | | Functionality | Stop Codon | Defective RSS |
| IGHV1-2 | P | ● | | IGHV1-3*03 | P | ● | | IGHV1-1 | P | ● | |
| IGHV1-3 | P | ● | | IGHV2-1*02 | ORF | | ● | IGHV1-11 | P | ● | |
| IGHV1-4 | P | ● | | IGHV3-1 | P | ● | | IGHV1-14 | P | ● | ● |
| IGHV3-3 | ORF | | ● | IGHV3-5*01 | P | ● | | IGHV3-3*02 | P | ● | |
| IGHV3-4 | P | ● | ● | IGHV4-4 | P | ● | | IGHV3-3*04 | P | ● | |
| IGHV3-14 | P | ● | | IGHV(III)-1 | P | ● | | IGHV3-16*01 | P | ● | |
| IGHV3-15 | ORF | | ● | | | | | IGHV3-16*02 | P | ● | |
| IGHV3-17 | P | ● | | | | | | IGHV3-17 | ORF | | ● |
| IGHV3-22 | P | ● | | | | | | IGHV4-1 | P | ● | |
| IGHV3-24 | P | ● | | | | | | IGHV4-3*04 | P | ● | |
| IGHV3-25 | ORF | | ● | | | | | IGHV4-3*05 | P | ● | |
| IGHV4-1 | P | ● | ● | | | | | IGHV9-3 | P | ● | |
| IGHV4-2 | P | ● | | | | | | IGHV9-4 | P | ● | |
| IGHV4-5 | P | ● | | | | | | IGHV9-5 | P | ● | |
| IGHV7-1 | P | ● | | | | | | IGHV9-6 | P | ● | ● |
| IGHV9-1 | P | ● | | | | | | IGHV9-7 | P | ● | |
| IGHV9-3 | P | ● | | | | | | IGHV9-8 | P | ● | |
| IGHV(II)-1 | P | ● | | | | | | IGHV9-9 | P | ● | ● |
| IGHV(II)-2 | P | ● | | | | | | IGHV10-2 | P | ● | |
| IGHV(II)- | P | ● | | | | | | IGHV11-3 | P | ● | ● |

| | | | | | | | |
|---|---|---|---|---|---|---|---|
| 3 | | | | | | | |
| IGHV(III)-1 | P | ● | | IGHV11-5 | P | ● | |
| IGHV(III)-2 | P | ● | | IGHV11-6 | P | ● | ● |
| IGHV(III)-3 | P | ● | ● | IGHV11-9 | P | ● | |
| | | | | IGHV(II)-1 | P | ● | |
| | | | | IGHV(II)-2*01 | P | ● | |
| | | | | IGHV(II)-2*02 | P | ● | |
| | | | | IGHV(II)-3 | P | ● | ● |
| | | | | IGHV(II)-4 | P | ● | ● |
| | | | | IGHV(III)-1 | P | ● | |
| | | | | IGHV(III)-2*01 | P | ● | |
| | | | | IGHV(III)-2*02 | P | ● | |
| | | | | IGHV(III)-3 | P | ● | ● |
| | | | | IGHV(III)-4 | P | ● | |

Table S4 IGH CDR3 HTS data sheet of human and mouse.

| Sample | IGH | | Subclasses | | | |
|---|---|---|---|---|---|---|
| | Productive | Clonotype | Name | Productive | Clonotype | Clonotype/Productive |
| M1 | 2,724,205 | 374,307 | IGG | 322,460 | 36,705 | 11.38% |
| | | | IGA | 639,297 | 71,378 | 11.17% |
| | | | IGM | 1,762,448 | 266,224 | 15.11% |
| M2 | 3,133,607 | 543,367 | IGG | 368,040 | 52,336 | 14.22% |
| | | | IGA | 725,174 | 103,608 | 14.29% |
| | | | IGM | 2,040,393 | 387,423 | 18.99% |
| M3 | 1,528,623 | 194,248 | IGG | 192,903 | 20,497 | 10.63% |
| | | | IGA | 324,177 | 38,988 | 12.03% |
| | | | IGM | 1,011,543 | 134,763 | 13.32% |
| H1 | 413,204 | 28,955 | IGG | 23,626 | 2,560 | 10.84% |
| | | | IGA | 218,596 | 9,793 | 4.48% |
| | | | IGM | 170,982 | 16,601 | 9.71% |
| H2 | 329,736 | 26,111 | IGG | 17,625 | 2,205 | 12.51% |
| | | | IGA | 207,771 | 8,683 | 4.18% |
| | | | IGM | 104,340 | 15,223 | 14.59% |
| H3 | 275,755 | 22,595 | IGG | 19,984 | 2,524 | 12.63% |
| | | | IGA | 154,748 | 8,635 | 5.58% |
| | | | IGM | 101,023 | 11,437 | 11.32% |

Table S5 Determination of Cytb genotypes of Rhinolophus affinis.

| Sample | Cytb Sequence | Identity | Score | Result |
|---|---|---|---|---|
| Bat1 | AGATTCAAAGGAACGCATTTCGTCGATCTACCAGCCCCCTCAAGTATTTCCTCCTGATGAAACTTCG<br>GATCCCTCCTAGGGGTCTGCCTTGCTGTACAAATTATTACAGGCCTTTTCCTAGCTATACACTACACA<br>TCAGACACCGCCACAGCCTTCTACTCTGTAACCCATATCTGCCGAGACGTCAACTACGGCTGAGTCC<br>TACGCTACCTCCATGCCAACGGAGCCTCCATATTCTTTATCTGCCTGTTCCTACACGTAGGACGAGGG<br>ATCTACTATGGCTCCTATACATTTTCAGAAACATGAAACATCGGAATTATCCTCCTCTTCGCCGTCAT<br>AGCCACAGCATTTATGGGCTATGTACTTCCATGAGGCCAAATATCCTTCTGAGGGGCAACAGTCATT<br>ACAAACCTCCTCTCAGCCATCCCCTATGTAGGAACAACCCTAGTAGAATGAGTCTGAGGAGGATTCT<br>CAGTAGACAAGGCCACACTCACCCGATTCTTCGCCTGACTTCCTCCCA | 99.57% | 850 | Rhinolophus affinis |
| Bat2 | TAAACGGACGCATTCGTCGATCTACCAGCCCCCTCAAGTATTTCCTCCTGATGAAACTTCGGATCCCT<br>CCTGGGGGTCTGCCTTGCTGTACAAATTATTACAGGCCTTTTCCTAGCTATACACTACACATCAGACA<br>CCGCCACAGCCTTCTACCCCGTAACCCATATCTGCCGAGACGTTAACTACGGCTGAGTCCTACGCTA<br>CCTCCATGCCAACGGAGCCTCCATATTCTTTATCTGCCTGTTCCTACACGTAGGACGAGGGATCTACT<br>ATGGCTCCTATACATTTTCAGAAACATGAAACATCGGAATTATCCTCCTCTTCGCCGTCATAGCCACA<br>GCATTTATGGGCTATGTACTTCCATGAGGCCAAATATCCTTCTGAGGGGCAACAGTCATCACAAACC<br>TCCTCTCAGCCATCCCCTATGTAGGAACAACCCTAGTAGAATGGGTCTGAGGAGGATTCTCAGTAAA<br>CAAAGCCACACTCACCCGATTCTTCGCCTGACTTTCCTCCAA | 99.10% | 798 | Rhinolophus affinis |
| Bat3 | CTAGGTCATGACGCATTCGTCGATCTACCAGCCCCCTCAAGTATTTCCTCCTGATGAAACTTCGGATC<br>CCTCCTAGGGGTCTGCCTTGCTGTACAAATTATTACAGGCCTTTTCCTAGCTATACACTACACATCAG<br>ACACCGCCACAGCCTTCTACTCTGTAACCCATATCTGCCGAGACGTCAACTACGGCTGAGTCCTACG<br>CTACCTCCATGCCAACGGAGCCTCCATATTCTTTATCTGCCTGTTCCTACACGTAGGACGAGGAATCT<br>ACTATGGCTCCTATACATTTTCAGAAACATGAAACATCGGAATTATTCTCCTCTTCGCCGTCATAGCC<br>ACAGCATTTATGGGCTATGTACTTCCATGAGGCCAAATATCCTTCTGAGGGGCAACAGTCATCACAA<br>ACCTCCTCTCAGCCGTCCCCTATGTAGGAACAACCCTAGTAGAATGGGTCTGAGGAGGATTCTCAGT<br>AGACAAGGCCACACTCACCCGATTCTTCGCCTGACCTTCCTCCCA | 99.38% | 874 | Rhinolophus affinis |

Figure S1 Amino acid sequence analysis of IGHV gene in three species of bats.

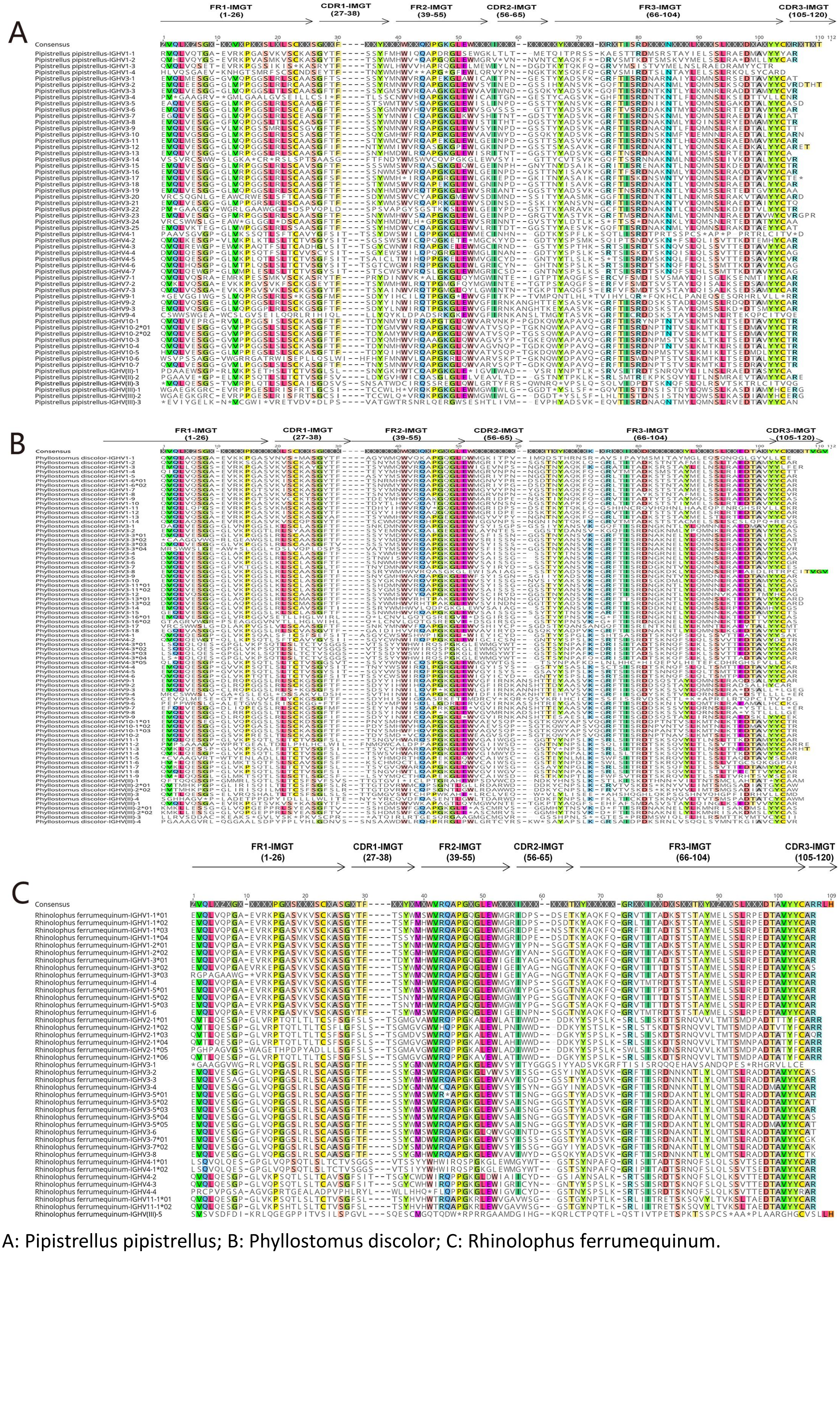


A: Pipistrellus pipistrellus; B: Phyllostomus discolor; C: Rhinolophus ferrumequinum.

Figure S2 Comparative analysis of IGHV genes in three bat species and other species.

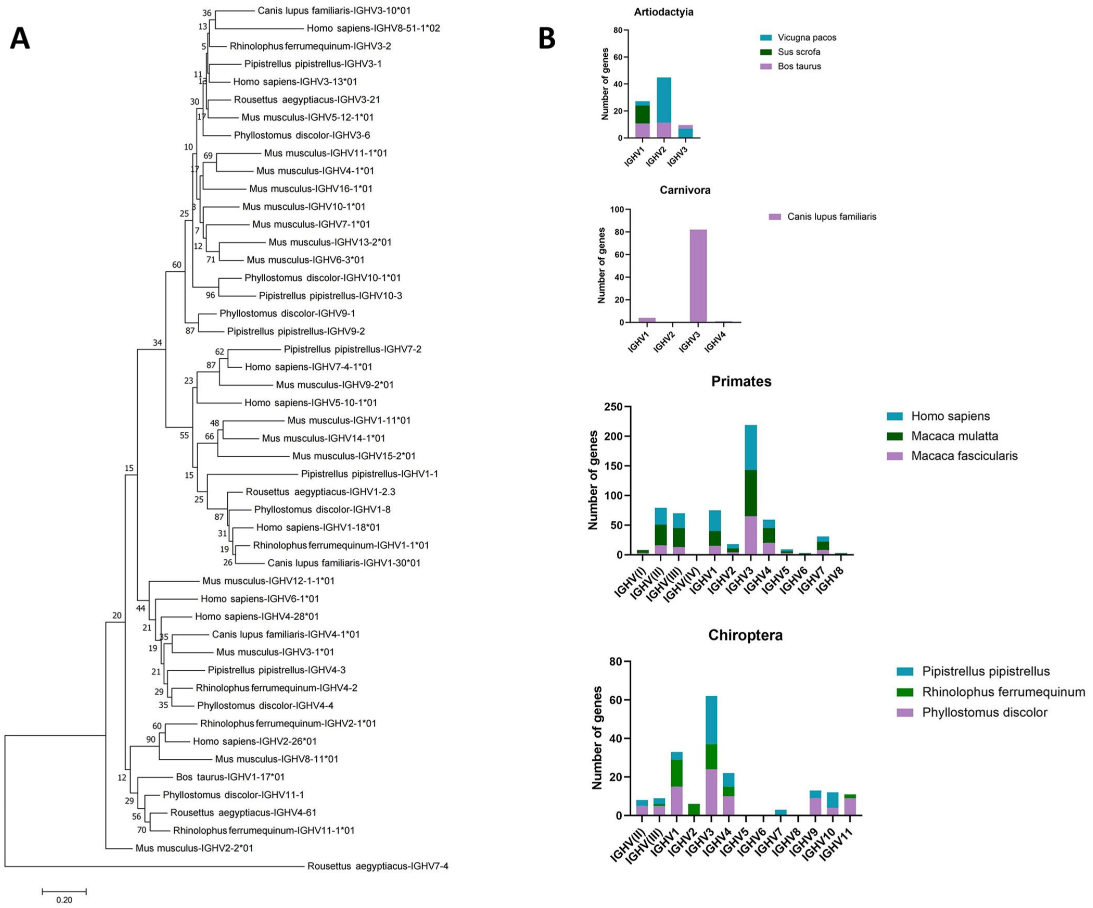


A: comparative analysis of IGHV gene phylogenetic tree of three species of bat, human, mouse, dog and cow;B: comparison of the number of IGHV gene families in chiroptera, carnivores, primates and artiodactyla.

Figure S3 Comparative analysis of the results of IGG quality control primers and experimental primers in Rhinolophus affinis.

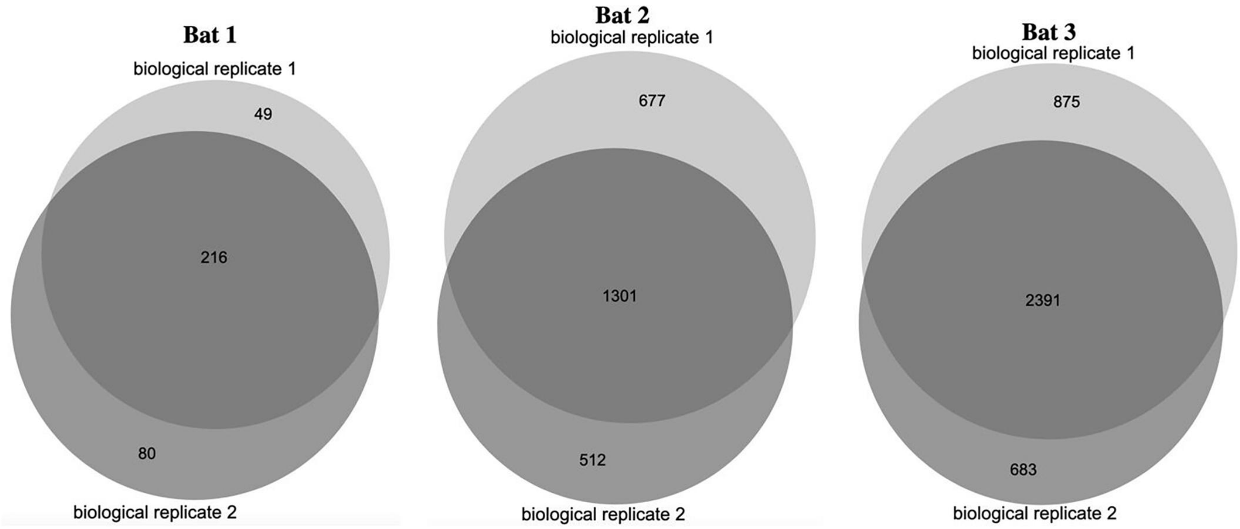

Figure S4 IGH CDR3 HTS data analysis of network sharing Human and Mouse.

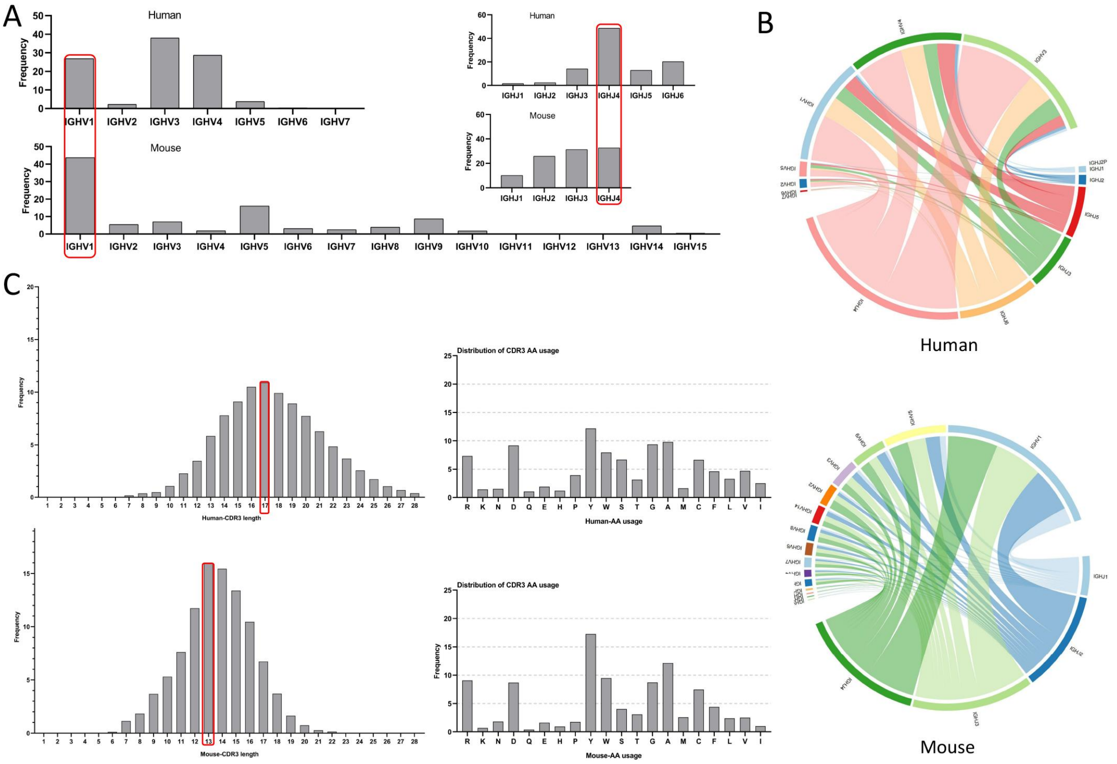


A: V and J usage analysis; B: V and J pairing analysis; C: Analysis of IGH CDR3 length and AA usage.

Figure S5 Analysis of V and J usage of Rhinolophus affinis, Human and Mouse IGM and IGE CDR3 group of Rhinolophus affinis and Mouse.

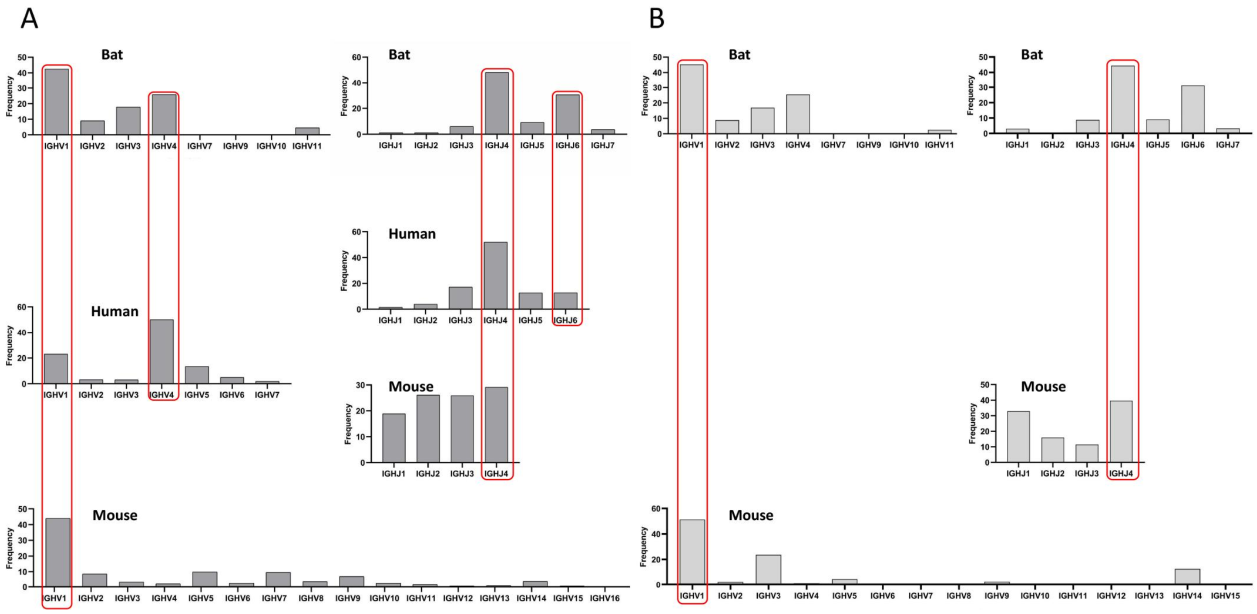


A: Analysis of V and J access of IGM CDR3 repertoire; B: Analysis of V and J access of IGE CDR3 repertoire.

Figure S6 V and J pairing of IGH CDR3 repertoire of Rhinolophus affinis, Human and Mouse.

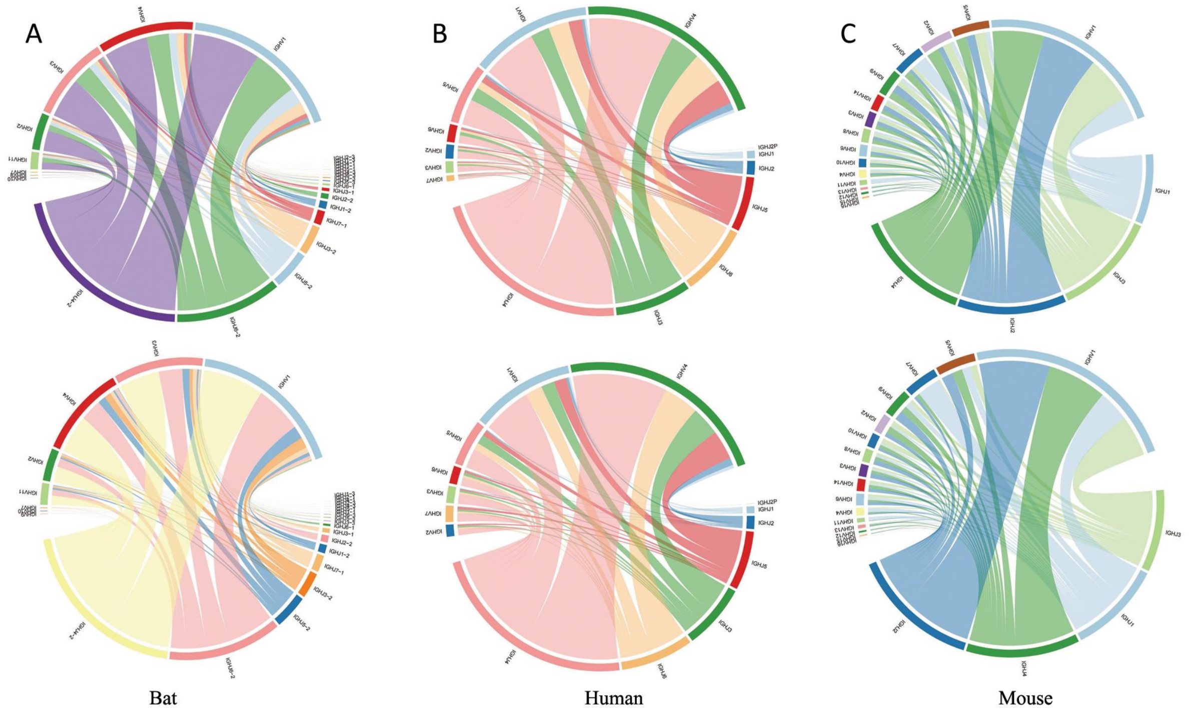


A: IGH CDR3 repertoire V and J pairing of B2 and B3; B: IGH CDR3 repertoire V and J pairing of H2 and H3; C: IGH CDR3 repertoire V and J pairing of M2 and M3.

Figure S7 Top clonal and relative abundance analysis, overlap analysis of IGH CDR3 subspecies (IGA, IGM, IGG) in Rhinolophus affinis, Human, and Mouse.

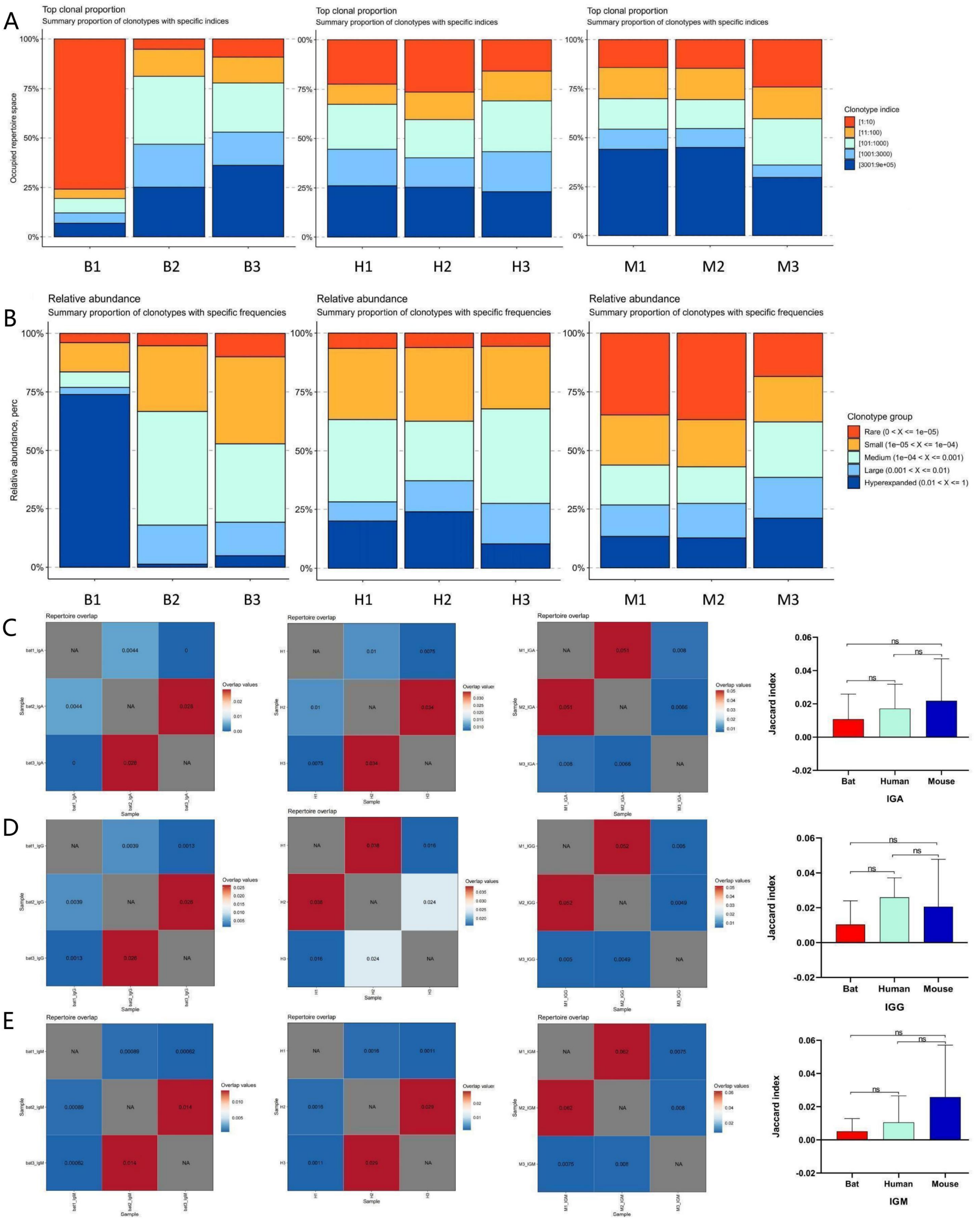


A: Top clonal analysis;B: Relative abundance analysis;C: Jaccard index of IGA;D: Jaccard index of IGG; E: Jaccard index of IGM.( ns: P>0.05; *: P<0.05; **: P<0.01; ***: P<0.001.)

Figure S8 Comparative analysis of IGHCDR3 total motif among Rhinolophus affinis, Human and Mouse.

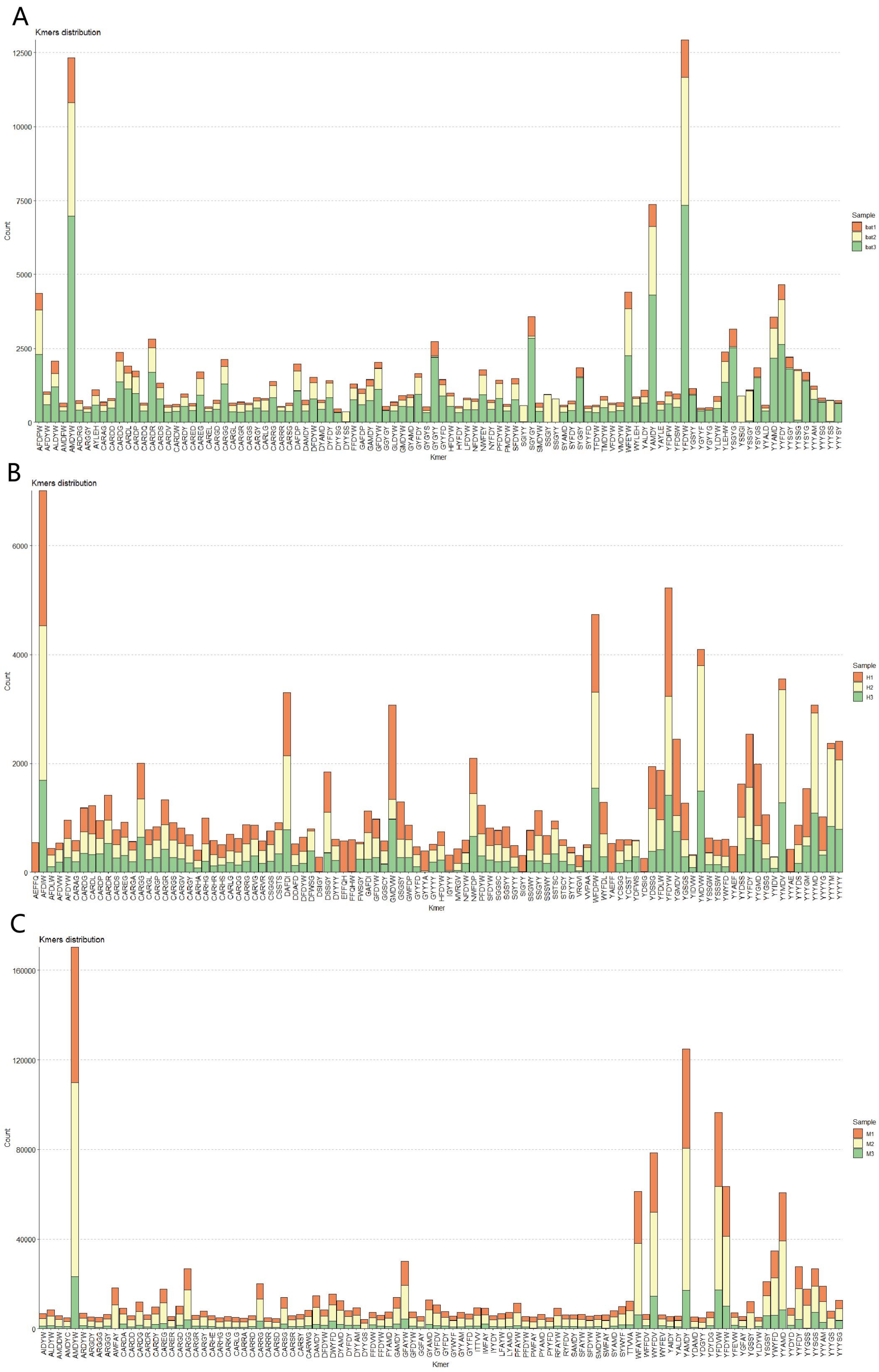


A: Bat;B: Human;C: Mouse

Figure S9 Rhinolophus affinis,Human,Mouse IGH CDR3 subclass moitf with the highest frequency of 10.

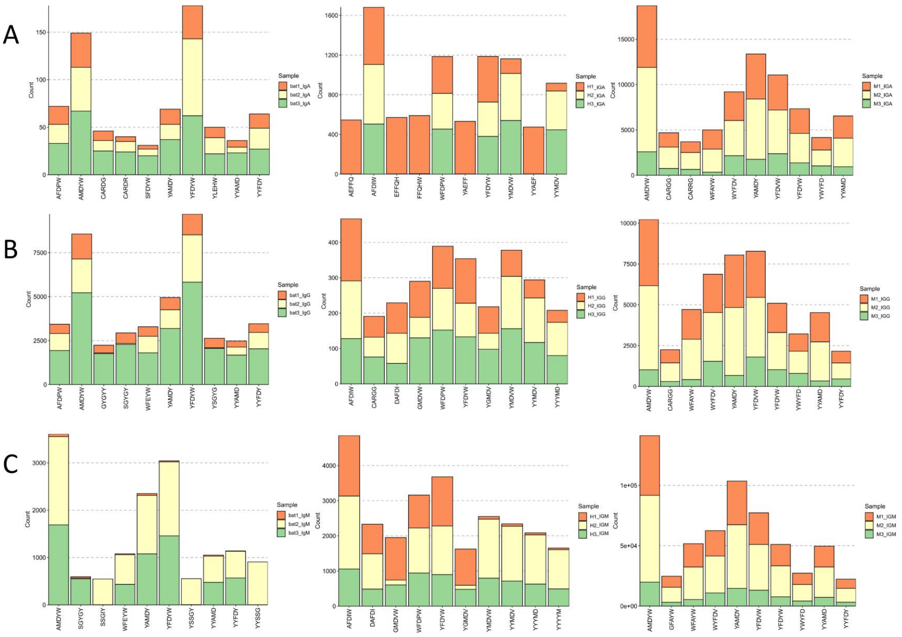

| Number | IGA | | |
|---|---|---|---|
| | Bat | Human | Mouse |
| | YFDYW | AFDIW | AMDYW |
| | AMDYW | YFDYW | YAMDY |
| | AFDPW | WFDPW | YFDVW |
| | YAMDY | YMDVW | WYFDV |
| | YYFDY | YYMDV | YFDYW |
| | YLEHW | FFQHW | YYAMD |
| | CARDG | EFFQH | WFAYW |
| | CARDR | AEFFQ | CARGG |
| | YYAMD | YAEFF | YWYFD |
| | SFDYW | YYAEF | CARRG |

| Number | IGG | | |
|---|---|---|---|
| | Bat | Human | Mouse |
| | YFDYW | AFDIW | AMDYW |
| | AMDYW | WFDPW | YFDVW |
| | YAMDY | YMDVW | YAMDY |
| | YYFDY | YFDYW | WYFDV |
| | AFDPW | YYMDV | YFDYW |
| | WFEYW | GMDVW | WFAYW |
| | SGYGY | DAFDI | YYAMD |
| | YSGYG | YGMDV | YWYFD |
| | YYAMD | YYYMD | CARGG |
| | GYGYY | CARGG | YYFDY |

| Number | IGM | | |
|---|---|---|---|
| | Bat | Human | Mouse |
| | AMDYW | AFDIW | AMDYW |
| | YFDYW | YFDYW | YAMDY |
| | YAMDY | WFDPW | YFDVW |
| | YYFDY | YMDVW | WYFDV |
| | WFEYW | YYMDV | WFAYW |
| | YYAMD | DAFDI | YFDYW |
| | YYSSG | YYYMD | YYAMD |
| | SGYGY | GMDVW | YWYFD |
| | YSSGY | YYYYM | GFAYW |
| | SSGIY | YGMDV | YYFDY |

A: IGA;B: IGG;C: IGM.

Figure S10 IGE overlap and motif analysis of Rhinolophus affinis and Mouse.

A

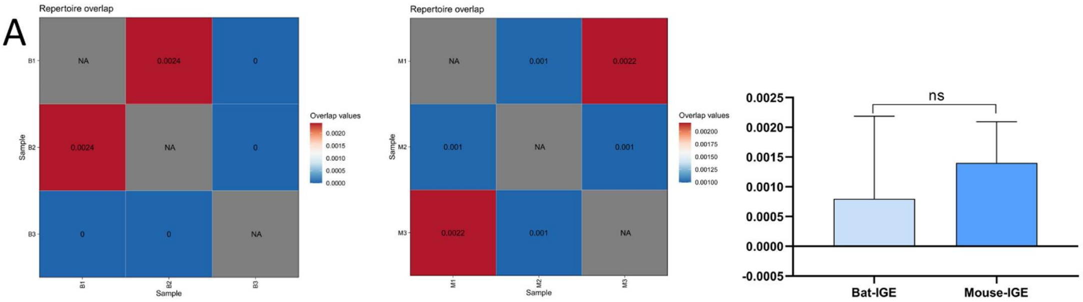


B

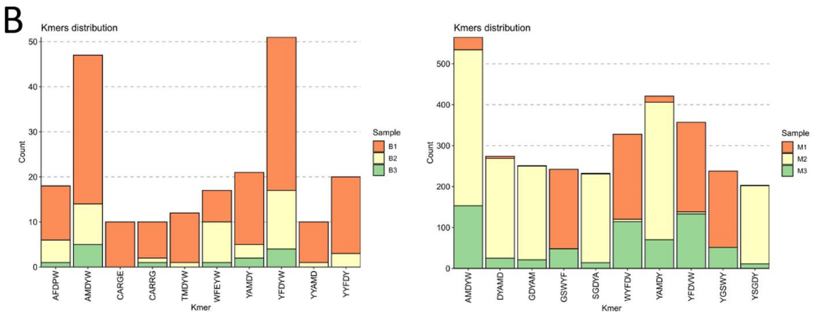


| Number | IGE | |
|---|---|---|
| | Bat | Mouse |
| ↓ | YFDYW | AMDYW |
| | AMDYW | YAMDY |
| | YAMDY | YFDVW |
| | YYFDY | WYFDV |
| | AFDPW | DYAMD |
| | WFEYW | GDYAM |
| | TMDYW | GSWYF |
| | CARGE | YGSWY |
| | CARRG | SGDYA |
| | YYAMD | YSGDY |

C

| Number | BAT | | | |
|---|---|---|---|---|
| | IGA | IGG | IGM | IGE |
| ↓ | YFDYW | YFDYW | AMDYW | YFDYW |
| | AMDYW | AMDYW | YFDYW | AMDYW |
| | AFDPW | YAMDY | YAMDY | YAMDY |
| | YAMDY | YYFDY | YYFDY | YYFDY |
| | YYFDY | AFDPW | WFEYW | AFDPW |
| | YLEHW | WFEYW | YYAMD | WFEYW |
| | CARDG | SGYGY | YYSSG | TMDYW |
| | CARDR | YSGYG | SGYGY | CARGE |
| | YYAMD | YYAMD | YSSGY | CARRG |
| | SFDYW | GYGYY | SSGIY | YYAMD |

A:IGE overlap analysis of Rhinolophus affinis and Mouse;

B:The 10 highest frequency motif in the IGE of Rhinolophus affinis and Mouse;

C:The 10 motif with the highest frequency among the four subspecies of Rhinolophus affinis.（Unpaired t test，nsP>0.05,*P<0.05,**P<0.01， ***P<0.001,****P<0.0001）

Figure S11 Comparative analysis of nucleotide sequences of IGH-CH1 in Rhinolophus affinis

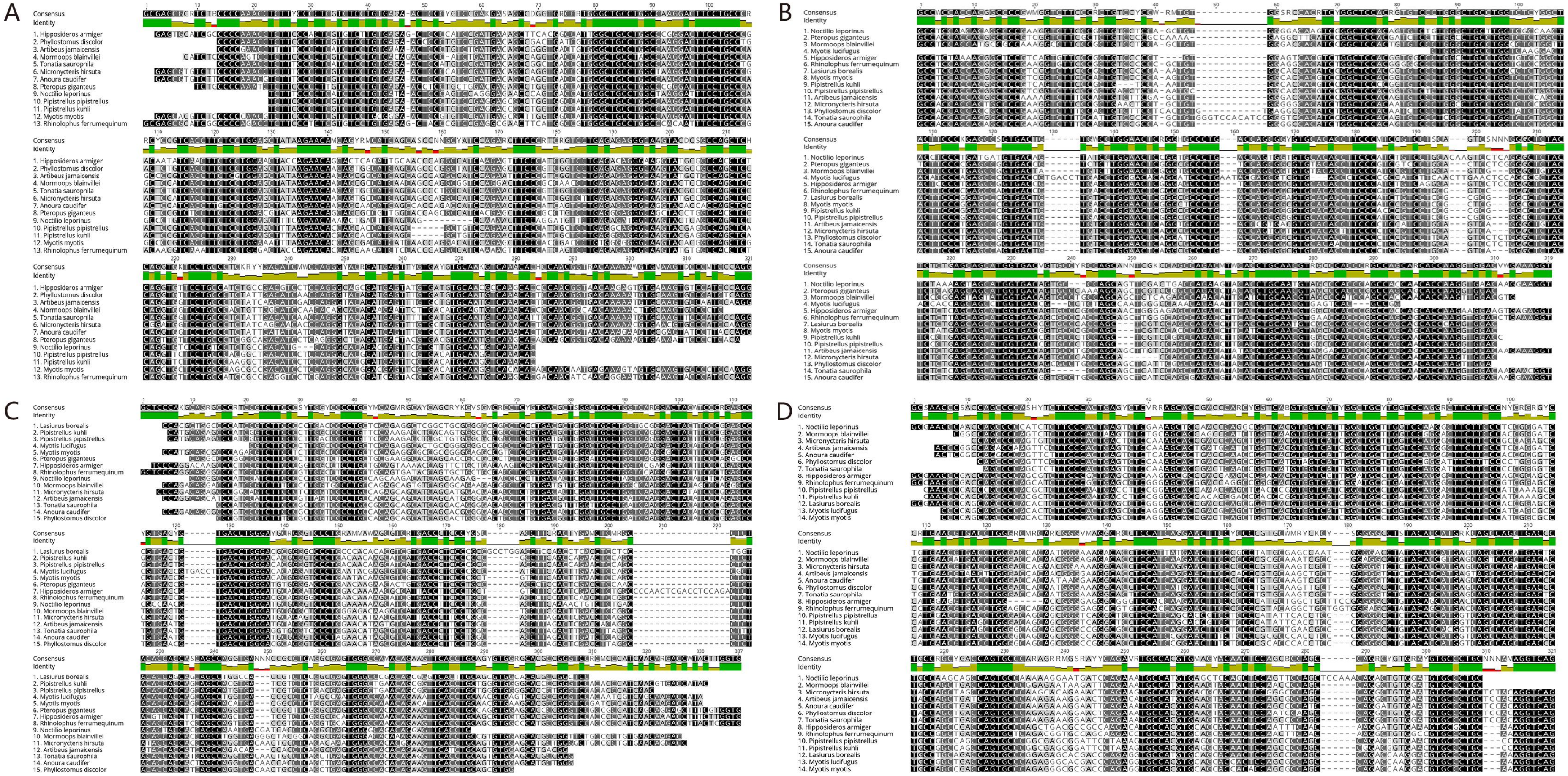

A: IGHM; B: IGHG; C: IGHE; D: IGHA.
